# Metabolic sensing in AgRP regulates sucrose preference and dopamine release in the nucleus accumbens

**DOI:** 10.1101/2023.12.14.571788

**Authors:** Alex Reichenbach, Harry Dempsey, Zane B. Andrews

## Abstract

Hunger increases the motivation for calorie consumption, often at the expense of low taste appeal. However, the neural mechanisms integrating calorie-sensing with increased motivation for calorie consumption remain unknown. Agouti-related peptide neurons in the arcuate nucleus of the hypothalamus sense hunger, and the ingestion of caloric solutions promote dopamine release in the absence of sweet taste perception. Therefore, we hypothesized that metabolic-sensing of hunger by AgRP neurons would be essential to promote dopamine release in the nucleus accumbens in response to caloric, but not non-caloric solutions. Moreover, we examined whether metabolic sensing in AgRP neurons affected taste preference to bitter solutions under conditions of energy need. Here we show that impaired metabolic sensing in AgRP neurons attenuated nucleus accumbens dopamine release in response to sucrose, but not saccharin, consumption. Further, metabolic sensing in AgRP neurons was essential to distinguish nucleus accumbens dopamine response to sucrose consumption when compared with saccharin. Under conditions of hunger, metabolic sensing in AgRP neurons increased the preference of sucrose solutions laced with the bitter tastant, quinine, to ensure calorie consumption whereas mice with impaired metabolic sensing in AgRP neurons maintained a strong aversion to sucrose/quinine solutions despite ongoing hunger. In conclusion, we demonstrate normal metabolic sensing in AgRP neurons drives the preference for calorie consumption, primarily when needed, by engaging dopamine release in nucleus accumbens.

## Introduction

Hunger has a profound effect to enact behaviours associated with food-seeking and consumption. A core feature of this hunger-driven behavioural repertoire is the motivation to continue food-seeking until energy requirements are met. Hypothalamic Agouti-related peptide (AgRP) neurons, located in the arcuate nucleus (ARC), are key hunger-sensing neurons in the brain (Aponte et al., 2011; Krashes et al., 2011; Luquet et al., 2005), whereas dopamine neurons in the ventral tegmental area (VTA) play a critical role in motivation. Indeed, there are many parallels between AgRP and dopamine neurons with respect to food-seeking and motivation. For example, dopamine-deficient and AgRP ablated mice are both hypophagic and prone to starvation (Luquet et al., 2005; Szczypka et al., 1999) and the activation of AgRP or dopamine neurons increases motivated food-seeking (Aponte et al., 2011; Krashes et al., 2011; Roitman et al., 2004).

Hunger increases dopamine release in the nucleus accumbens in response to food or cues predicting food (Aitken et al., 2016; Mazzone et al., 2020; Wallace et al., 2020) highlighting the interaction between homeostatic and motivational systems. Chemogenetic activation of AgRP neurons increase nucleus accumbens dopamine release and VTA dopamine neural activity to food (Alhadeff et al., 2019; Mazzone et al., 2020), demonstrating that AgRP neurons relay information pertaining to homeostatic state to motivation pathways. Indeed, impaired metabolic sensing in AgRP neurons weakens the ability to transmit homeostatic information between AgRP neurons and dopamine pathways (Reichenbach et al., 2022). This impaired metabolic sensing in AgRP neurons not only reduced dopamine release to sucrose pellets, it also suppressed motivated food seeking during fasting.

Hunger not only increases motivation for food, but it also modifies liquid taste preference to ensure calorie intake. For example, fasted mice will consume more sucrose solution at a lower concentration or consume more sucrose solution adulterated with a bitter tastant (Fu et al., 2019) and this effect was replicated by the chemogenetic or optogenetic activation of AgRP neurons (Fu et al., 2019). Moreover, cues predicting sucrose consumption produces a greater phasic release of dopamine compared to cues predicting saccharin (McCutcheon et al., 2011), and mice lacking sweet taste receptors prefer sucrose over sucralose (de Araujo et al., 2008). These results imply that AgRP neurons promote the consumption of caloric solutions, which are inherently rewarding and elicit a dopamine response independent from taste. In our current studies, we examined whether metabolic sensing in AgRP neurons distinguished caloric from non-caloric solutions by regulating dopamine release in the nucleus accumbens. Moreover, we examined whether metabolic sensing in AgRP neurons affected taste preference to bitter solutions under conditions of energy need. To do this, we used the deletion of carnitine acetyltransferase (*Crat*) in AgRP as a previously validated model of impaired metabolic sensing under conditions of energy need (Reichenbach et al., 2022; Reichenbach et al., 2018a; Reichenbach et al., 2018b; Reichenbach et al., 2018c).

## Results

### Metabolic sensing in AgRP neurons facilitates dopamine release to sucrose intake

To test if metabolic sensing in AgRP neurons helps to distinguish caloric sucrose solutions from sweet but non-caloric saccharin solutions by regulating dopamine release, we exposed AgRP Crat WT or KO mice to 4% sucrose and 0.1% saccharin for 20 minutes while recording nucleus accumbens dopamine release using GRAB-DA (Sun et al., 2018) (Fig 1 A,B). All mice were naïve to caloric and non-caloric sweetened solutions to ensure an untrained dopamine response. In response to sucrose consumption, AgRP Crat KO mice had significantly lower dopamine release in the nucleus accumbens compared to WT mice (Fig 1 C,D). In response to saccharin consumption, however, the increase in dopamine release was not different between genotype (Fig 1 E,F). Furthermore, when comparing the effect of sucrose and saccharin on dopamine release in the nucleus accumbens (Fig 1 G,I), we observed greater persistence of the dopamine signal after sucrose consumption compared to saccharin in WT mice (Fig 1 H), but not KO mice (Fig 1 J). Thus, metabolic sensing in AgRP neurons was necessary for the appropriate dopamine response to sweet caloric solutions but not sweet non-caloric solutions, indicating that AgRP neurons transmit caloric information into a persistent dopamine signal within the nucleus accumbens.

**Figure 1:**
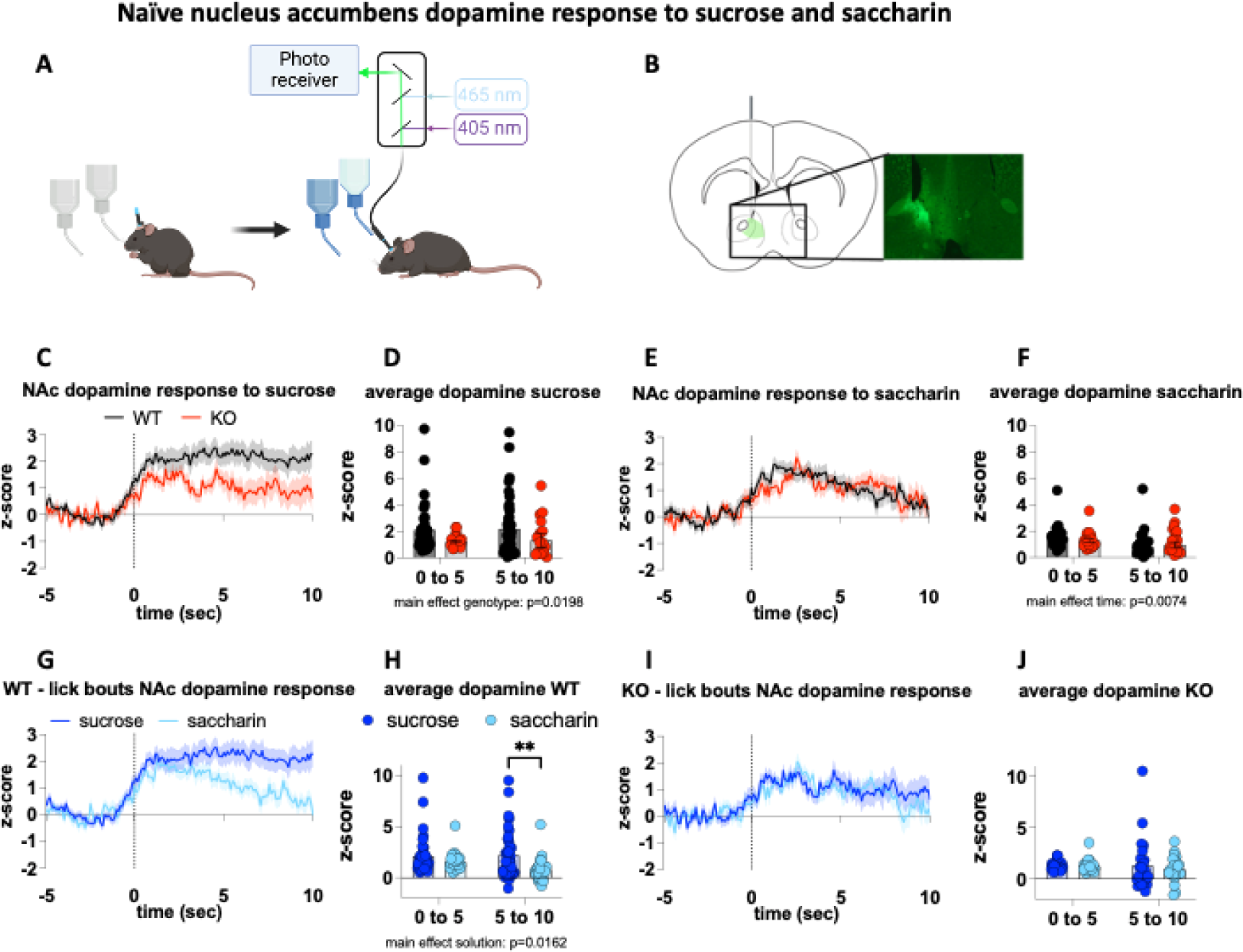
Impaired metabolic sensing in AgRP neurons affects NAc dopamine response to sucrose but not saccharin solutions. Naïve WT and AgRP Crat KO mice expressing GRAB Da sensor in the nucleus accumbens were exposed to 4% sucrose and 0.1% saccharin solutions and dopamine response during lick bouts recorded (A,B). Dopamine response to sucrose (C) and saccharin bouts (E) and average z-score during response (D+F). Comparison of dopamine signal during sucrose and saccharin bouts and average z-score of early (0-5 sec) and late (5-10 sec) response in WT (G,H) and KO (I,J).

Based on persistent dopamine release in response to a caloric sucrose solution in WT mice, we hypothesised that WT and KO mice would show significant differences in sucrose preference under fed and fasted conditions. To test this idea, mice were placed in BioDaq cages to allow for precise measurement of fluid intake over time (Fig 2 A). Initial 2 bottle comparison between water and saccharin (Fig 2 B,C) reveal a clear preference for the sweet taste of saccharin for both genotypes, indicating impaired metabolic sensing in AgRP neurons does not affect taste preference for sweet solutions, an observation that is supported by similar dopamine responses to saccharin in WT and KO mice (Fig 1 E). Given the choice between 4% sucrose and 0.1% saccharin, mice consume more sucrose than saccharin irrespective of genotype (Fig 2 D,E) and the sweet taste of solutions increases the fluid intake compared to the average water intake (Fig 2J). However, the increased need for calories during fasting elevates the preference for sucrose more in WT than in KO (Fig 2 F-I). Thus, metabolic sensing in AgRP neurons potentiates the shift towards sucrose preference during fasting to restore energy balance.

**Figure 2:**
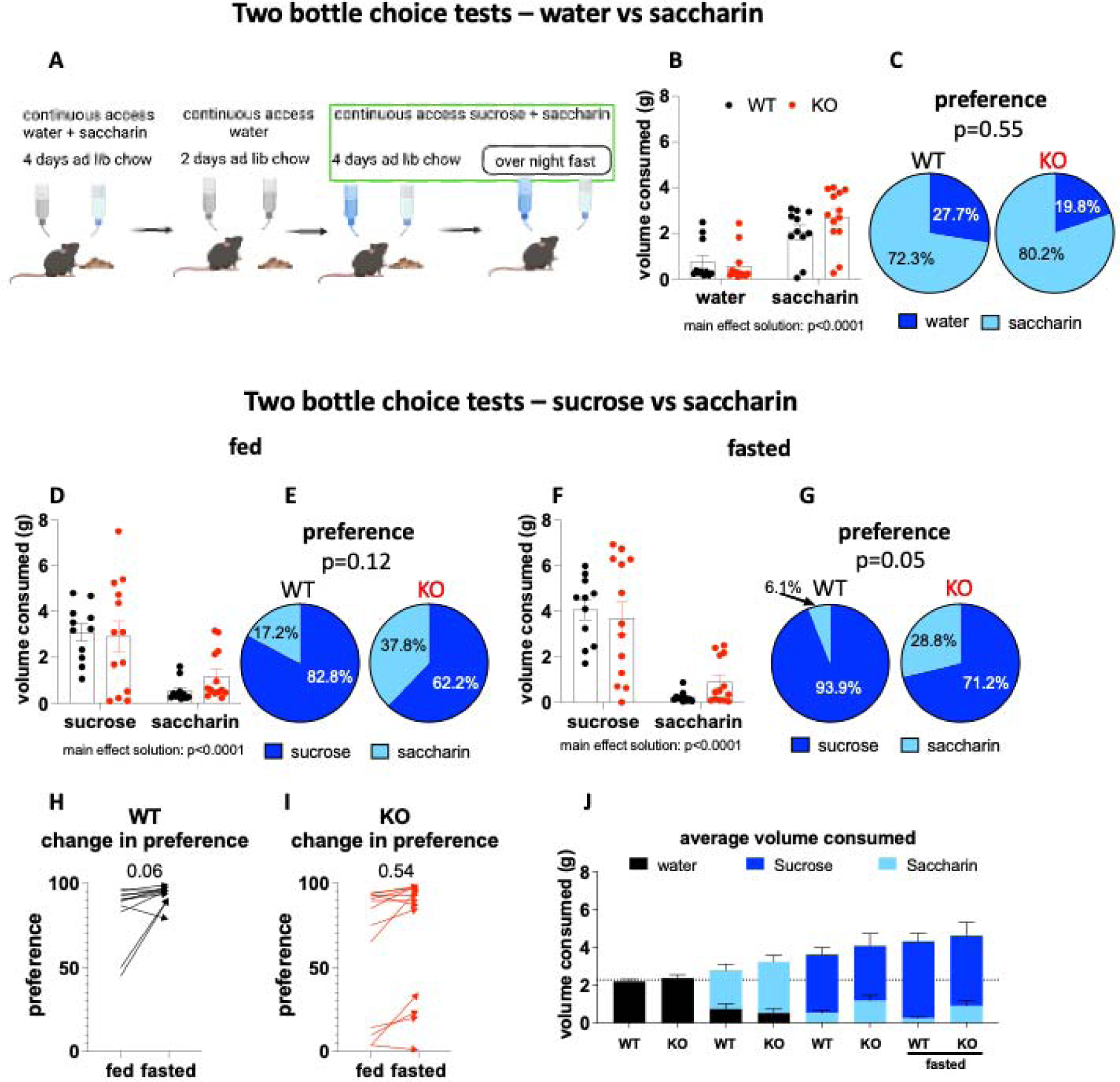
impaired metabolic sensing in AgRP neurons impedes sucrose preference. In a 2-bottle choice paradigm, male and female mice had ad libitum access to chow for 4 days followed by an overnight fast. Position of bottles were swapped daily and volume consumed overnight recorded. Bottles were washed after each choice pair and mice had 2 days of water in both bottles (A). Pie charts compare preference between genotypes for saccharin (light blue) and comparator solution as indicated (dark blue). After acclimation to the BioDaq cages, mice received choice of water and 0.1% saccharin (B+C), followed by 4% sucrose and 0.1% saccharin with ad libitum access to chow (D+E) or without access to chow (F+G). Fasting induced change in sucrose preference in WT (H) and KO (I) and average total (saccharin plus comparator solution) fluid consumed per day (J). Data +/-SEM, two-way ANOVA with Tukey’s post hoc analysis and unpaired student’s t-test; a, significant at p<0.05.

Bitter tastes such as quinine or denatonium are often used as adulterants to create taste aversions and suppress intake. However, both fasting and AgRP neuronal stimulation drive greater consumption of sucrose despite the presence of bitter tastes (Fu et al., 2019), implying AgRP neurons monitor energy state and prioritise caloric intake over taste. To test whether metabolic sensing in AgRP neurons is important in this process, we performed sucrose/quinine - saccharin preference tests (Fig 3 A). Pairing the caloric solution with an unpalatable taste shifts the preference to saccharin in KO, with WT mice showing no preference (Fig 3 B,C). Strikingly, an overnight fast increases the preference for the quinine-laced sucrose solution in WT, but not in KO mice (Fig 3 D,E), resulting in a significant increase in preference from fed to fasted in WT mice (Fig 3 F) but no change in preference in KO mice (Fig 3 G). Thus, AgRP neurons require intact metabolic-sensing capabilities to increase caloric consumption by increasing sucrose preference despite the bitter quinine taste.

**Figure 3:**
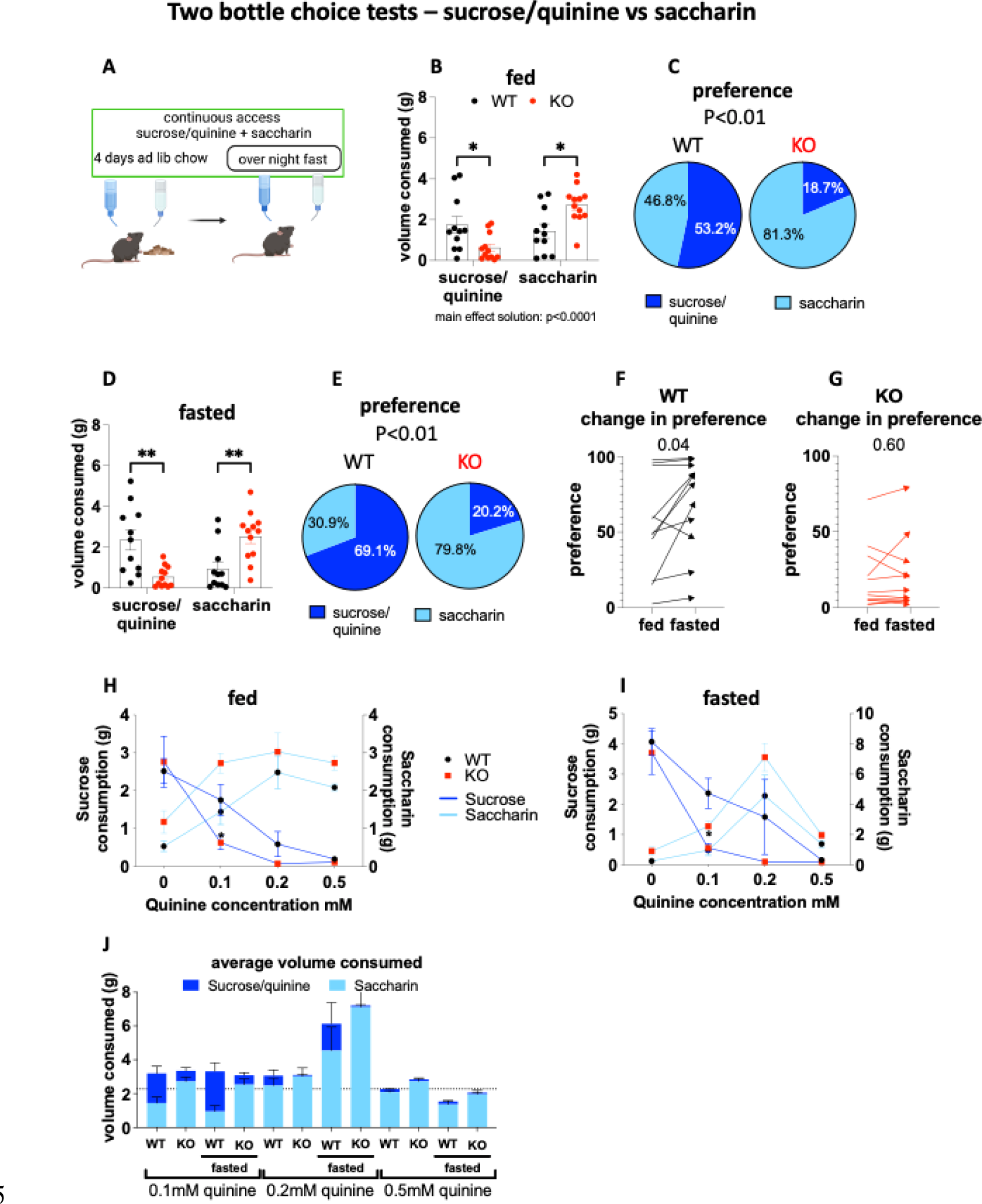
Metabolic sensing in AgRP neurons enables choice of calories over taste during fasting. Same 2 bottle choice paradigm as in Figure 2 but with 0.1% saccharin and 4% sucrose laced with 0.1mM Quinine (A). Fluids consumed (B) and preference (C) during ad libitum access to chow and during fasting (D+E). Fasting induced changes in sucrose/quinine preference in WT (F) and KO (G). Dose response curve for quinine concentrations in 4% sucrose solution from different cohorts of mice (n=5-14) ad libitum fed (H) and without access to chow (I) and combined saccharin and sucrose/quinine average volume consumed (J). Data +/- SEM, two-way ANOVA with Tukey’s post hoc analysis and unpaired student’s t-test; a, significant at p<0.05.

The dramatic increase in saccharin preference over sucrose/quinine solutions in KO mice highlights potentially greater sensitivity to bitter tastes or stronger taste avoidance. To examine this further, we examined 4% sucrose consumption laced with 0, 0.1, 0.2, 0.5 mM quinine in fed and fasted mice. Increasing the quinine concentrations in the sucrose solution gradually reduced the volume consumed with 0.5mM quinine being too unpalatable for mice to drink (Fig 3 H-J). In fed mice, lacing sucrose with 0.1mM quinine did not reduce the volume consumed in WT compared to 4% sucrose but it significantly reduced the consumption in KO mice so that they consumed significantly less compared to WT. Importantly, inducing a metabolic need state by removing food during the 16 hours access increased consumption in WT but not KO mice, resulting in a significant main effect of genotype for overall consumption (Fig 3 I). This suggests that metabolic sensing in AgRP neurons balances the caloric value of sucrose with the tolerance for aversive taste. Collectively, these results show that metabolic sensing in AgRP neurons during fasting increases caloric sucrose intake by increasing the tolerance for quinine, which is used to devalue the taste of sucrose.

## Discussion

In these studies, we assessed how metabolic sensing in AgRP neurons affected sucrose preference and NAc dopamine release. To model impaired metabolic sensing in AgRP neurons we deleted Crat from AgRP neurons and previously demonstrated aberrant glucose sensing using *ex vivo* electrophysiological recordings or *in vivo* AgRP GCaMP6 recordings (Reichenbach et al., 2022). This metabolic sensing defect is also supported by 1) a proteomic screen in which AgRP neurons lacking Crat show a number of changes in mitochondrial and metabolic pathways (Reichenbach et al., 2018b); and 2) impaired peripheral metabolism in response to refeeding after fasting or prolonged food restriction (Reichenbach et al., 2018a; Reichenbach et al., 2018b; Reichenbach et al., 2018c).

With this model, we demonstrated that impaired metabolic sensing in AgRP neurons attenuated NAc dopamine release in response to sucrose, but not saccharin, consumption. Further, WT mice displayed a greater NAc dopamine response to sucrose when compared with saccharin, in line with previous reports showing that cues predicting sucrose produce larger increases in dopamine release compared to those predicting saccharin (McCutcheon et al., 2011). As there were no genotype differences in NAc dopamine release to saccharin, our results suggest metabolic sensing in AgRP neurons is essential to transmit neural information about caloric content, but not sweet taste, to dopamine terminals in the NAc.

Previous studies showed mice lacking sweet taste receptors could still increase striatal dopamine release and form a preference for caloric (sucrose) solutions compared to non-caloric sucralose solutions (de Araujo et al., 2008). We suggest metabolic sensing in AgRP neurons may be the essential step to form sucrose preference in the absence of sweet taste perception, as reported (de Araujo et al., 2008). Based on this logic, the absence of caloric-processing may shift preferences towards sweet solutions independent of calorie content or need. In line with this, the preference for saccharin over sucrose laced with quinine didn’t change in fed or fasted KO mice. Thus, the inability to discern caloric and non-caloric sweetened solutions meant that choice behaviour was guided mainly by taste. A similar observation was reported after ablating AgRP neurons in neonatal mice, as feeding relied more on palatability and dopamine tone (Denis et al., 2015).

As AgRP neurons are critical for sensing hunger, our results suggest the effects of hunger on NAc dopamine release (Wallace et al., 2020) in response to food (Mazzone et al., 2020) or cues predicting food (Aitken et al., 2016) are driven by AgRP neurons. This assumption is further supported by studies showing the activation of AgRP neurons increases NAc dopamine release (Alhadeff et al., 2019; Mazzone et al., 2020), a response that is required for AgRP neurons to drive appropriate food motivation during fasting (Reichenbach et al., 2022). Whether or not metabolic sensing in AgRP neurons affects VTA DA cell firing in a similar manner remains to be determined. In support of this view, chemogenetic activation of ARC NPY (AgRP) neurons elevates VTA dopamine activity to food presentation (Mazzone et al., 2020). However, NAc dopamine release can occur independent from changes from VTA neural firing, a process thought to dissociate roles for dopamine in learning and motivation (Mohebi et al., 2019). Intriguingly, chemogenetic activation of AgRP neurons drives NAc dopamine release (Alhadeff et al., 2019; Mazzone et al., 2020) but reduces the number of active VTA dopamine neurons following intragastric Ensure infusion (Grove et al., 2022). Thus, it is possible that these opposing AgRP-driven changes at the NAc nerve terminal and VTA cell bodies mediate different behaviours, although this requires further experimental evidence.

Our results show that impaired metabolic sensing in AgRP neurons diminished sucrose preference, which is consistent with previous observations showing that chemogenetic AgRP activation and inhibition increased and decreased sucrose licking activity respectively (Fu et al., 2019). The inability of KO mice to increase their preference for sucrose in the fasted state, as occurs in WT mice, highlights the key role for AgRP neurons to guide behaviour based on appropriate metabolic sensing of energy need. Indeed, AgRP neurons encode caloric information into neural firing properties as caloric foods, but not non-caloric foods, produces larger and sustained inhibition of AgRP neuronal firing (Beutler et al., 2017; Chen et al., 2015; Su et al., 2017). Our previous studies demonstrate that impaired metabolic sensing in AgRP neurons affects encoding of caloric information in response to glucose and peanut butter (Reichenbach et al., 2022).

AgRP neurons, however, also affect taste preference since the activation of AgRP neurons increased licking rate, whereas AgRP inhibition decreased licking rate, for sucrose laced with a bitter taste (Fu et al., 2019). Our results showed a similar response in which fasting increased consumption of sucrose laced with quinine in WT but not KO mice. Indeed, KO mice showed a strong preference for saccharin, which was unaffected by fasting. These results suggest that taste preference is mediated in part by the ability of hunger-sensing AgRP neurons to sense energy need, as observed in WT mice where caloric need caused by fasting increased sucrose/quinine consumption compared to fed mice. Although the influence of hunger on taste sensitivity has been well documented (Haase et al., 2009; Hanci and Altun, 2015), our results establish a key role for metabolic sensing in AgRP neurons.

Furthermore, saccharin-induced NAc dopamine release is similar in WT and KO mice, suggesting KO mice have a stronger taste aversion for sucrose/quinine consumption. Thus, impaired metabolic sensing in AgRP neurons may affect taste aversion, although future studies are required to explore this possibility. Although a dopamine signal of aversive taste occurs separate from gut-brain processing of caloric value (Stauffer et al., 2015), both require appropriate subsequent, and potentially overlapping, neural integration. In fact, gut-brain feedback carries both post-ingestive calorie information (Beutler et al., 2017; Su et al., 2017) and malaise signals (Zimmerman et al., 2023) and is one potential way that AgRP neurons could integrate caloric information with taste aversion. These post-ingestive malaise signals target regions with AgRP neural terminals, including the parabrachial nucleus and amygdala (Betley et al., 2013). Moreover, AgRP neurons dynamically encode the caloric value of nutritive signals in learned flavour nutrient pairings (Beutler et al., 2017; Nyema et al., 2023; Su et al., 2017) making them perfect candidates to mediate post-ingestive taste-nutrient associations.

In summary, metabolic sensing in AgRP is required to correctly transmit and transform AgRP hunger-signaling into NAc dopamine release after calorie consumption. Importantly metabolic sensing in AgRP neurons does not affect NAc dopamine release after sweet non-caloric saccharin consumption highlighting different mechanisms used to distinguish calorie need from sweet taste. Furthermore, metabolic sensing in AgRP neurons adjusts the preference to caloric-containing solutions under conditions of energy need, even at the expense of aversive bitter tastants. Overall, these studies demonstrate that hunger-sensing AgRP neurons drive the preference for calorie consumption, primary when needed, by engaging dopamine release in NAc. The abnormal function of this pathway may contribute to impaired feeding behaviour and lead to either under or over-eating, which characterise human conditions such as eating disorders and obesity.

## Methods

### Animals

Male and female mice were kept under standard laboratory conditions with free access to food (chow diet, catalog no. 8720610, Barastoc Stockfeeds, Victoria, Australia) and water at 23C in a 12-hr light/ dark cycle and were group-housed to prevent isolation stress unless otherwise stated. All mice were aged 8 weeks or older for experiments unless otherwise stated. *Agrp*-ires-cre mice were obtained from Jackson Laboratory AgRP^tm1(cre)Low/J^ (stock no. 012899; The Jackson Laboratory, Maine, USA). To delete Crat from AgRP neurons, *Agrp*-ires-cre mice were crossed with Crat^fl/fl^ mice (Randall Mynatt, Pennington Biomedical Research Center, LA, USA) (AgRP^cre/wt^::Crat^fl/fl^ mice; designated as KO). AgRP^wt/wt^::Crat^fl/fl^ littermate mice were designated as WT and used as controls. All experiments were conducted in compliance with the Monash University Animal Ethics Committee guidelines.

### Stereotaxic surgery and fiber implantation

Animals were anesthetised (2-3% isoflurane) and injected with Metacam (5 mg/kg) prior to placing into heated (37C) stereotaxic frame (Stoelting) and viral injections to target the nucleus accumbens (bregma 1.2mm, midline 0.5 mm. skull -4.8mm) were performed as previously described. Non-cre dependent dopamine sensor (YL10012-AAV9: AAV-hSyn-DA4.3 (Sun et al., 2018) was unilaterally injected for 6 min @25nl/min followed by 5 min rest before withdrawing the pulled glass pipette. Ferrule capped fibers (400µm core, NA 0.48 Doric, MF1.25 400/430-0.48) were implanted above the injection site and secured with dental cement (GBond, Japan). Mice had at least 2 weeks recovery before commencement of experiments.

### Fiber photometry

Fiber photometry experiments were performed using optical components from Doric lenses controlled by Tucker Davis Technologies fiber photometry processor RZ5P., Mice for fiber photometry experiments were housed in modified home cages with excess to 2 sipper bottles (filled with water) through a side wall. For experiments examining NAc dopamine release mice were tethered and acclimatized to the fiber cable in their home cage without access to bottles. During recording mice had access to sipper bottles with 0.1% saccharin and 4% sucrose and drinking bouts were time-stamped using a home cage sipper device ((Godynyuk et al., 2019)). Dopamine responses to 0.1% saccharin and 4% sucrose were measured in naïve mice during 20 minutes recording session. Contact with sipper bottles were confirmed through video analysis (Openscope, Tucker Davis Technologies) and only bouts longer than one second and with an immediate change in dopamine signal of at least 1 z-score from baseline were included for analysis. Custom python codes were used to filter and analyse fiber photometry data, which is available here https://github.com/Andrews-Lab/Fiber_photometry_analysis.

### Two bottle choice tests

Male and female mice were single housed in BioDaq feeding cages with ad libitum access to chow and two water bottles during 3 days of acclimatization. For water vs saccharin consumption, one bottle was filled with 0.1% saccharin and position of bottles swapped daily 2 hours before onset of dark phase. After 4 days ad libitum feeding, mice were overnight fasted (5pm-9am) with only access to drink bottles followed by 2 days wash out period with ad lib access to water and chow. In the same manner as above, mice were presented with choice of 4% sucrose and 0.1% saccharin solutions or 4% sucrose laced with quinine HCl . (concentrations for different cohorts ranging from 0.1 to 0.5mM) and 0.1% saccharin to measure consumption in fed and fasted states.

### Statistical analysis

Statistical analyses were performed using GraphPad Prism for MacOS X. Data are represented as mean ± SEM. Two-way ANOVAs with post hoc tests were used to determine statistical significance. A two-tailed Student’s unpaired t-test (all statistical information is supplied in a detailed supplementary table) was used when comparing genotype only. p < 0.05 was considered statistically significant.

## Supporting information

Supplementary Statistical Table

