## Supplementary Statistical Table for "Metabolic sensing in AgRP regulates sucrose preference and dopamine release in the nucleus accumbens"

### Appendix 1. Statistical test details and results for all analyses

| Figure | Statistical test | Group n | Main analysis result | Significant post-hoc multiple comparisons |
|---|---|---|---|---|
| **1C** | 2-way RM ANOVA | WT (n =39 trails from 5 mice)<br>KO (n=21 trails from 3 mice) | Genotype $F$(1, 58) = 6.374 ; **p = 0.0143**<br>Time $F$(5.753, 333.7) = 12.08; **p < 0.0001**<br>Genotype x Time $F$(152, 8816) = 1.829; **p < 0.0001** | |
| **1D** | 2-way RM ANOVA | | Genotype $F$(1, 122) =5.576; **p = 0.0198**<br>Time $F$(1, 122) = 0.03645; p =0.8489<br>Genotype x Time $F$(1, 122) = 4.183e-007; p = 0.9995 | |
| **1E** | 2-way RM ANOVA | WT (n =26 trails from 4 mice)<br>KO (n=23 trails from 4 mice) | Genotype $F$(1, 47) = 0.07805 ; p = 0.7812<br>Time $F$(12.72, 598.0) = 12.35; **p < 0.0001**<br>Genotype x Time $F$(152, 7144) = 1.307; **p = 0.0072** | |
| **1F** | 2-way RM ANOVA | | Genotype $F$(1, 94) =0.1038; p = 0.7480<br>Time $F$(1, 94) = 7.503; **p =0.0074**<br>Genotype x Time $F$(1, 94) = 1.193; p = 0.2775 | |
| **1G** | 2-way RM ANOVA | sucrose (n =39 trails from 5 mice)<br>saccharin (n=26 trails from 4 mice) | Solution $F$(1, 63) = 6.299 ; p = 0.0147<br>Time $F$(6.268, 394.9) = 17.07; **p < 0.0001**<br>Solution x Time $F$(152, 9576) = 2.792; **p < 0.0001** | Sidak's multiple comparison's test:<br>Sucrose – saccharin row 153: **p = 0.0319** |
| **1H** | 2-way RM ANOVA | | Solution $F$(1, 63) = 6.103 ; **p = 0.0162**<br>Time $F$(1, 63) = 2.744; p = 0.1026<br>Solution x Time $F$(1, 63) = 3.959; p = 0.0510 | Sidak's multiple comparison's test:<br>Sucrose – saccharin 5 to 10 seconds: **p = 0.0045** |
| **1I** | 2-way RM ANOVA | sucrose (n =21 trails from 3 mice)<br>saccharin (n=23 trails from 4 mice) | Solution $F$(1, 38) = 0.4160 ; p = 0.5228<br>Time $F$(9.171, 348.5) = 6.645; **p < 0.0001**<br>Solution x Time $F$(152, 5776) = 0.8630; p = 0.8853 | |
| **1J** | 2-way RM ANOVA | | Solution $F$(1, 63) = 0.2370 ; p = 0.6288<br>Time $F$(1, 63) = 0.2403; p = 0.6264<br>Solution x Time $F$(1, 63) = 0.5666; p = 0.4555 | |

| Figure | Statistical test | Group n | Main analysis result | post-hoc multiple comparisons |
|---|---|---|---|---|
| **2B** | 2-way ANOVA | WT (n =11)<br>KO (n=13) | Genotype $F$(1, 44) = 0.6142 ; p = 0.4374<br>Solution $F$(1, 44) = 36.14; **p < 0.0001**<br>Genotype x Solution $F$(1, 44) = 2.290; p = 0.1374 | |
| **2C** | 2-way ANOVA | | Genotype $F$(1, 44) = 5.577e-017 ; p > 0.9999<br>Solution $F$(1, 44) = 32.60; **p < 0.0001**<br>Genotype x Solution $F$(1, 44) = 0.7282; p = 0.3981 | Uncorrected Fisher's LSD:<br>Water WT vs KO: p = 0.5493<br>Saccharin WT vs KO: p = 0.5493<br>WT water vs saccharin: **p = 0.0019**<br>KO water vs saccharin: **p < 0.0001** |
| **2D** | 2-way ANOVA | | Genotype $F$(1, 44) = 0.2727 ; p = 0.6041<br>Solution $F$(1, 44) = 23.91; **p < 0.0001**<br>Genotype x Solution $F$(1, 44) = 0.8881; p = 0.3511 | |
| **2E** | 2-way ANOVA | | Genotype $F$(1, 44) = 1.243e-015 ; p > 0.9999<br>Solution $F$(1, 44) = 32.60; **p < 0.0001**<br>Genotype x Solution $F$(1, 44) = 5.224; **p = 0.0271** | Uncorrected Fisher's LSD:<br>sucrose WT vs KO: p = 0.1132<br>Saccharin WT vs KO: p = 0.1132<br>WT sucrose vs saccharin: **p < 0.0001**<br>KO sucrose vs saccharin: p = 0.0525 |
| **2F** | 2-way ANOVA | | Genotype $F$(1, 44) = 0.1023 ; p = 0.7506<br>Solution $F$(1, 44) = 50.56; **p < 0.0001**<br>Genotype x Solution $F$(1, 44) = 1.200; p = 0.2793 | |
| **2G** | 2-way ANOVA | | Genotype $F$(1, 44) = 6.872e-015 ; p > 0.9999<br>Solution $F$(1, 44) = 68.80; **p < 0.0001**<br>Genotype x Solution $F$(1, 44) = 1.200; **p = 0.0059** | Uncorrected Fisher's LSD:<br>sucrose WT vs KO: **p = 0.0466**<br>Saccharin WT vs KO: **p = 0.0466**<br>WT sucrose vs saccharin: **p < 0.0001**<br>KO sucrose vs saccharin: **p = 0.0002** |
| **2H** | Unpaired t-test | n = 11 | t = 1.942; p = 0.0663 | |
| **2I** | Unpaired t-test | n = 13 | t = 0.6106; p = 0.5472 | |
| **Figure** | **Statistical test** | **Group n** | **Main analysis result** | **Significant post-hoc multiple comparisons** |
| **3B** | 2-way ANOVA | WT (n =11)<br>KO (n=12) | Genotype $F$(1, 42) = 0.06637 ; p = 0.7980<br>Solution $F$(1, 42) = 8.513; **p = 0.0056** | Sidak's multiple comparison's test:<br>Sucrose/quinine WT vs KO: **p = 0.0259**<br>Saccharin WT vs KO: **p = 0.0101** |

| | | | | |
|---|---|---|---|---|
| | | | Genotype x Solution $F(1, 42) = 15.41$; **p = 0.0003** | |
| **3C** | 2-way ANOVA | | Genotype $F(1, 42) = 1.068e-016$ ; p > 0.9999<br>Solution $F(1, 42) = 12.17$; **p = 0.0012**<br>Genotype x Solution $F(1, 42) = 18.13$; **p = 0.0001** | Uncorrected Fisher's LSD:<br>Sucrose/quinine WT vs KO: **p = 0.0042**<br>Saccharin WT vs KO: **p = 0.0042**<br>WT sucrose/quinine vs saccharin: p = 0.5869<br>KO sucrose/quinine vs saccharin: **p < 0.0001** |
| **3D** | 2-way ANOVA | | Genotype $F(1, 42) = 0.06637$ ; p = 0.7980<br>Solution $F(1, 42) = 8.513$; **p = 0.0056**<br>Genotype x Solution $F(1, 42) = 23.55$; **p = 0.0003** | Sidak's multiple comparison's test:<br>Sucrose/quinine WT vs KO: **p = 0.0014**<br>Saccharin WT vs KO: **p = 0.0052** |
| **3E** | 2-way ANOVA | | Genotype $F(1, 42) = 0.1067$ ; p = 0.7456<br>Solution $F(1, 42) = 0.6697$; p = 0.4178<br>Genotype x Solution $F(1, 42) = 18.13$; **p < 0.0001** | Uncorrected Fisher's LSD:<br>Sucrose/quinine WT vs KO: **p = 0.0042**<br>Saccharin WT vs KO: **p = 0.0042**<br>WT sucrose/quinine vs saccharin: p = 0.5869<br>KO sucrose/quinine vs saccharin: **p < 0.0001** |
| **3F** | Unpaired t-test | n = 11 | t = 2.399; p = 0.0374 | |
| **3G** | Unpaired t-test | n = 12 | t = 0.4803; p = 0.6404 | |
| **3H sucrose** | 2-way ANOVA | WT:<br>0 Q: n = 10<br>0.1 Q n = 11<br>0.2 Q n = 7<br>0.5 Q = 10<br><br>KO:<br>0 Q: n = 14<br>0.1 Q n = 12<br>0.2 Q n = 5<br>0.5 Q = 14 | Genotype $F(1, 75) = 1.240$ ; p = 0.3012<br>Quinine concentration $F(3, 75) = 17.09$; **p < 0.0001**<br>Genotype x Quinine concentration $F(3, 75) = 1.568$; p = 0.2144 | Tukey's multiple comparisons test<br>0.1: WT vs KO: **p = 0.0373**<br>WT: 0 vs 0.2: **p = 0.0150**<br>0 vs 0.5: **p = 0.0006**<br>0.1 vs 0.5: **p = 0.0313**<br>KO: 0 vs 0.1: **p = 0.0003**<br>0 vs 0.2: **p = 0.0007**<br>0 vs 0.5: **p < 0.0001** |
| **3H saccharin** | 2-way ANOVA | | Genotype $F(1, 75) = 14.65$ ; **p = 0.0003**<br>Quinine concentration $F(3, 75) = 17.80$; **p < 0.0001**<br>Genotype x Quinine concentration $F(3, 75) = 0.7731$; p = 0.5127 | Tukey's multiple comparisons test<br>0.1: WT vs KO: **p = 0.0008**<br>WT: 0 vs 0.2: **p = 0.0001**<br>0 vs 0.5: **p = 0.0008**<br>KO: 0 vs 0.1: **p = 0.0002**<br>0 vs 0.2: **p = 0.0008**<br>0 vs 0.5: **p < 0.0001** |
| **3I sucrose** | 2-way ANOVA | | Genotype $F(1, 63) = 4.306$ ; **p = 0.0421**<br>Quinine concentration $F(3, 63) = 15.81$; **p < 0.0001** | Tukey's multiple comparisons test<br>0.1: WT vs KO: **p = 0.0168**<br>WT: 0 vs 0.2: **p = 0.0251** |

| | | | | |
|---|---|---|---|---|
| | | | Genotype x Quinine concentration *F*(3, 63) = 1.009; p = 0.3946 | 0 vs 0.5: **p = 0.0007**<br>KO: 0 vs 0.1: **p = 0.0002**<br>0 vs 0.2: **p = 0.0014**<br>0 vs 0.5: **p = 0.0003** |
| **3I saccharin** | 2-way ANOVA | | Genotype *F*(1, 63) = 12.55 ; **p = 0.0008**<br>Quinine concentration *F*(3, 63) = 32.93; **p < 0.0001**<br>Genotype x Quinine concentration *F*(3, 63) = 1.332; p = 0.2720 | Tukey's multiple comparisons test<br>0.1: WT vs KO: **p = 0.0140**<br>0.2: WT vs KO: **p = 0.0049**<br>WT: 0 vs 0.2: **p < 0.0001**<br>0.1 vs 0.2: **p < 0.0001**<br>0.2 vs 0.5: **p = 0.0033**<br>KO: 0 vs 0.1: **p = 0.0410**<br>0 vs 0.2: **p < 0.0001**<br>0.1 vs 0.2: **p < 0.0001**<br>0.2 vs 0.5: **p < 0.0001** |
